# Enhanced Neural Plasticity of the Primary Visual Cortex in Visual Snow Syndrome: Evidence from MEG Gamma Oscillations

**DOI:** 10.1101/2025.02.12.637794

**Authors:** E.V. Orekhova, S.M Naumova, T.S. Obukhova, A.M. Plieva, A.O. Prokofiev, A.V. Petrokovskaia, A.R. Artemenko, T.A. Stroganova

## Abstract

Visual Snow Syndrome (VSS) is a neurological disorder characterized by persistent visual disturbances and associated symptoms. Although the neural basis of VSS remains poorly understood, it may involve increased neuronal excitability and/or altered neuroplasticity in the visual cortex, which could, in turn, affect visual gamma oscillations. An altered excitation-inhibition (E-I) balance is hypothesized to alter the modulation of gamma power and frequency by stimulation intensity, while maladaptive neuroplasticity may impact time-dependent changes in gamma power during repeated stimulation.

To investigate potential alterations in E-I balance and neuroplasticity in VSS, we magnetoencephalography to record visual gamma oscillations in 26 VSS patients and 27 healthy controls. Participants were exposed to repeatedly presented high-contrast annular gratings, which were either static or drifting at varying speeds to systematically manipulate stimulation intensity. We also measured heart rate variability (HRV) during rest and repetitive visual stimulation to explore the relationship between time-dependent gamma changes and parasympathetic activation, which is known to promote activity-dependent plasticity.

Our results showed no significant group differences in gamma power or frequency, nor in their modulation by drift rate, suggesting that the excitation-inhibition (E-I) balance in the primary visual cortex remains largely intact in VSS. Both groups exhibited an initial brief decrease in gamma power followed by a sustained linear increase with stimulus repetition, likely reflecting activity-dependent plasticity. HRV parameters were comparable across groups, with the parasympathetic-sympathetic balance index correlating with repetition-related increase in gamma power, further supporting the link between time-dependent gamma changes and neuroplasticity. Notably, VSS patients exhibited a steeper repetition-related increase in gamma power, indicating atypically heightened activity-dependent plasticity in this group.

These findings provide the first experimental evidence suggesting that altered activity-dependent neuroplasticity plays a role in the pathophysiology of VSS. Furthermore, they identify repetition-related increases in gamma power as a potential biomarker of aberrant neuroplasticity, offering novel insights into VSS pathophysiology and potential avenues for targeted therapeutic interventions.

## Introduction

Visual Snow Syndrome (VSS) is a neurological disorder marked by the continuous presence of small, flickering dots throughout the entire visual field, resembling snowflakes or noise on an improperly tuned television. In addition to this visual ‘snow’, individuals with VSS often experience other visual phenomena, such as increased sensitivity to light (photophobia), difficulty seeing at night (nyctalopia), a persistence of images or their traces after stimuli disappearance (palinopsia), etc., and non-visual symptoms (e.g. migraine, tinnitus, trouble concentrating, fatigue, and anxiety)^1–4^.

While exact causes of VSS are unknown, it is widely hypothesized to be related to neural hyperexcitability in visual cortical networks^5,6^. It remains unclear, however, to what extent this hyperexcitability is linked to a deficit in inhibition^7^. Additionally, the degree of involvement of the primary visual cortex (V1) and secondary visual cortical areas in VSS is still uncertain. While many studies emphasize the role of areas downstream of V1, such as the lingual gyrus^8–10^ or higher-order supramodal associative areas^11^, there is also evidance for structural^12,13^ and functional^6,14–16^ abnormalities in the area V1, although the latter findings remain inconsistent. Detecting the earliest affected cortical level would enhance our understanding of the VSS mechanisms and, ultimately, aid in tailoring diagnosis and therapeutic intervention for the syndrome.

The role of neural excitability alternations in V1 in the pathophysiology of VSS may be elucidated by examining the properties of narrow-band visual gamma oscillations in patients with this disorder. This high-frequency oscillatory activity (40-90 Hz) is induced predomimantly in V1^17,18^ in response to stimuli of certain characteristics, such as high-contrast static gratings, especially those moving with a certain ‘optimal’ speed^19^, and can be reliably recorded non-invasively with MEG. Extensive animal research has shown that gamma oscillations are generated through recurrent interaction between cortical inhibitory and excitatory neurons, with significant imbalances in their activity influencing the frequency and power of gamma oscillations (e.g.^20–22^). The only MEG study of patients with VSS reported increased gamma power in response to static high-contrast gratings, which was interpreted as indicative of heightened neural excitability^14^. While this finding is intriguing and warrants independent replication, estimating MEG gamma power induced by a specific stimulus generally does not provide a reliable measure of excitation-inhibition (E-I) balance in V1 circuitry. One limitation stems from substantial inter-individual variability in this parameter, which can be influenced by factors unrelated to excitability, such as e.g., signal-to-noise ratio^23^ and cortical anatomy^18,24^— both of which may be altered in clinical populations. Another, more fundamental limitation stems from the dependence of gamma power on the activity of both excitatory and inhibitory neurons^25^. Animal studies has shown that, under different conditions, increased gamma synchronization may be caused by the activation of either excitatory^22^ or inhibitory^20^ neurons. As a result, the single-stimulus gamma response power may not excibit a simple linear relationship with shifts in the E-I balance in the visual cortex of patients with VSS.

Inferring the E-I balance based on the modulation of MEG gamma response power by increasing excitatory drive to the visual cortex presents a more promising approach. We previously demonstrated that increasing the drift rate of a visual grating—a proxy for excitatory drive— caused bell-shaped changes in gamma response power^19^. This non-linear modulation is likely due to the increasing excitation of inhibitory neurons, which initially enhance gamma oscillations but disrupt gamma synchronization at the excessively high level of excitatory drive^26,27^. A deficient inhibition is expected to alter the bell-shaped modulation curve, shifting the optimum for gamma generation toward higher values of excitatory drive. The relational nature of the gamma modulation curve overcomes the limitations of the single-stimulus approach, providing better sensitivity to abnormal E-I balance in V1 in neuropsychiatric disorders compared to absolute gamma response power^28,29^.

***The first aim*** of the present study was to assess the E-I balance in the early visual cortex of individuals with VSS by modulating the oscillatory gamma response through alternations in the drift rate of visual grating. We hypothesized that if the E-I ratio in patients with VSS is elevated due to inhibition deficit, this should be reflected in the atypical modulation of gamma response power by an increase in excitatory drive.

Beyond altered neural excitability, changes in neural plasticity have been proposed as potential pathophysiological mechanisms underlying both the symptoms and the functional and structural alterations in visual and extra-visual areas in VSS^30^. Neural plasticity refers to synaptic changes in the network-level interactions driven by prior neural activity, enaibling the neural network to adapt stimulus processing based on sensory experience^31^. While maladaptive synaptic plasticity has been clearly implicated in ‘phantom sensations’ in somatosensory (e.g., chronic pain) and auditory (e.g., tinnitus) domains^32–35^, its role in visual snow symptoms is unclear. To date, this hypothesis remains speculative, as there is a lack of experimental studies investigating neural plasticity in VSS.

Recent studies demonstrate that visual gamma oscillations recorded by MEG offer a means to investigate neuroplasticity induced by stimulus repetition in the early visual cortex (Peter, Stauch). Repeated exposure to the same visual stimuli is a fundamental aspect of visual experience that alters neural network properties^36–39^. While a repetition of an identical visual stimulus typically results in decreased neuronal firing rates^38,39^, and reduction of evoked responses in EEG/MEG^40–42^, it is also associated with an increase in gamma response power and frequency^38,41^. These repetition-related changes in gamma are stimulus-specific and can persist for tens of minutes after the exposure ends. Animal studies have demonstrated that the stimulus-specific increase in gamma power likely reflects Hebbian plasticity, which reshapes receptive fields in V1 for frequently exposed stimulus, thereby supporting implicit perceptual learning^37,38^. We hypothesized that a maladaptive increase in neuronal plasticity in patients with VSS may lead to inappropriate modifications in V1 circuitry, contributing to abnormal sensations in the absence of an actual visual stimulus. Our ***second aim*** was to investigate, for the first time, potential alterations in neural plasticity in VSS by examining repetition-related changes in visual gamma parameters in this population compared to healthy controls. We were also interested in examining the concomitant repetition-related changes in MEG evoked responses, given their proposed link to impaired habituation in patients with VSS^6^ and their likely co-occurance with maladaptive neuronal plasticity, as indexed by time-dependent changes in gamma synchronization.

Hebbian plasticity in cortical circuits is enhanced during states of arousal and attention, processes associated with a release of neuromodulators^43,44^ facilitating plasticity^45^. Fluctuations in attentional states^46^ and implicit learning^47^ are in turn linked to activation of the parasympathetic system. In individuals with VSS, parasympathetic activity may differ from that of controls due to the frequent presence of comorbid psychiatric symptoms such as anxiety and depression^48^. Therefore, our ***third aim*** was to investigate the role of autonomic regulation in V1 plasticity in both neurotypical individuals and those with VSS. To this end, we assessed heart rate variability (HRV) in relation to repetition-related changes in visual gamma responses. HRV served as a measure of the autonomic balance between the sympathetic and parasympathetic systems^49,50^, providing insights into the potential impact of this balance on neuroplastic processes in VSS.

In summary, the primary focus of the current study was to test the hypotheses of an elevated excitation-inhibition ratio and altered neuroplasticity in the primary visual cortex of patients with VSS. To address these hypotheses, we investigated changes in gamma oscillations related to variations in excitatory input to the visual cortex, as well as their time-dependent modulation in response to repeated stimuli.

## 2. Materials and methods

### 2.1. Participants

Twenty-six patients were recruited from an online VSS community and through clinic referrals. All patients underwent neurological examination, during which detailed information about their neurological and visual symptoms was collected. The examination was conducted by a board-certified neurologist who is also the author of this study (AA). Thirteen subjects from the patient group reported visual snow (VS) from early childhood (approximately before age 10), while the other 13 developed VS after age 14. Subjects who developed VSS due to substance abuse were excluded from the study. Nearly all participants (24 out of 26) were medication-free. Two participants were taking antidepressants: one on venlafaxine (75 mg daily) and another on escitalopram (5 mg daily), and both had been on their respective medications for more than three months.

We also recruited 27 neurologically healthy control participants from the community, matched for sex and age. The characteristics of the subjects are presented in Table 1. All subjects were asked to complete a Russian version of the Visual Discomfort Scale^51^, which assesses unpleasant somatic and perceptual side effects of pattern viewing. They were also asked to complete the Russian version of the State-Trait Anxiety Inventory^52^.

**Table 1.** Characteristics of the participants.

|  | VSS | Control | t, p |
| --- | --- | --- | --- |
| N | 26 (14 males) | 27 (16 males) |  |
| Age | 26.7 +/- 5.6 | 26.9 +/- 4.2 | n.s. |
| Migraine | 20 (77%) | 0% |  |
| Migraine with aura | 7 (27%) | 0% |  |
| Tinnitus | 14 (54%) | 0% |  |
| STAI (all life)* | 51.4 +/- 9.5 | 40.1 +/- 9.6 | t(51) = 4.3, p=0.00008 |
| Visual Discomfort score** | 25.9 +/- 13.1 | 8.5 +/-7.9 | t(50) = 5.8 p<0.00001 |

\* Information was not available for one VSS and one control participants
\*\* Information was not available for two VS and one control participants

The local ethics committee approved the study and all subjects gave written informed consent.

All but one patient with VSS rated a range of their visual symptoms on a 5-point Likert scale ( 1 – absence of the symptom, 2 – very rare, 3 – rare, 4 – often, 5 – all the time). Table 2 lists the number of subjects reported ‘4’ or ‘5’ on this scale for the symptoms most frequently associated with VSS.

**Table 2.** Visual phenomena in patients with VSS Visual.

| Visual symptoms | N (%) |
| --- | --- |
| Palinopsia (I): afterimages | 15 (58%) |
| Palinopsia (II): trailing of moving objects | 10 (38%) |
| Enhanced entopic phenomena | 19 (73%) |
| Photophobia | 19 (73%) |
| Impaired night vision (Nyctalopia) | 15 (58%) |

The majority of VSS subjects also had other visual symptoms, such as flashes of light, swirls or clouds with eyes closed, or other forms of unusual visual phenomena.

### 2.2. Experimental paradigm

The recording session began with two resting conditions: a 5-minute period with eyes closed, followed by a 5-minute period with eyes open. The ‘eyes closed’ condition was not analyzed in this study, whereas data from the ‘eyes open’ condition were used to estimate resting-state heart rate variability.

The visual task (Figure 1A) was presented immediately after the resting state. Participants watched a sequence of large high-contrast annular gratings (18°, full contrast, spatial frequency 1.66 cycles per degree) that either drifted toward the center at one of four velocities (0.6°/s, 1.2°/s, 3.6°/s, or 6.0°/s) or remained static. All trials started with a fixed 1.2 s prestimulus interval during which a fixation cross was presented in the center of the visual field. The presentation time for each stimulus ranged randomly from 1.2 to 1.6 seconds. After this period, the stimulus disappeared. Participants were instructed to press a button as soon as this happened. The new trial started immediately after the button press. If the button was not pressed within 1 second after the stimulus disappearance, the trial was considered a ‘missed trial’, and the text “You’re late!” was shown to the participant for 1.0 second. If the subject pressed the button earlier than 150 ms after stimulus disappearance, the trial was considered a ‘commission error trial’. To reduce fatigue and boredom, participants were shown short (3–6 s) cartoon animations after every 5-10 gratings. For each participant, 90 gratings of each type were presented.

**Figure 1.**
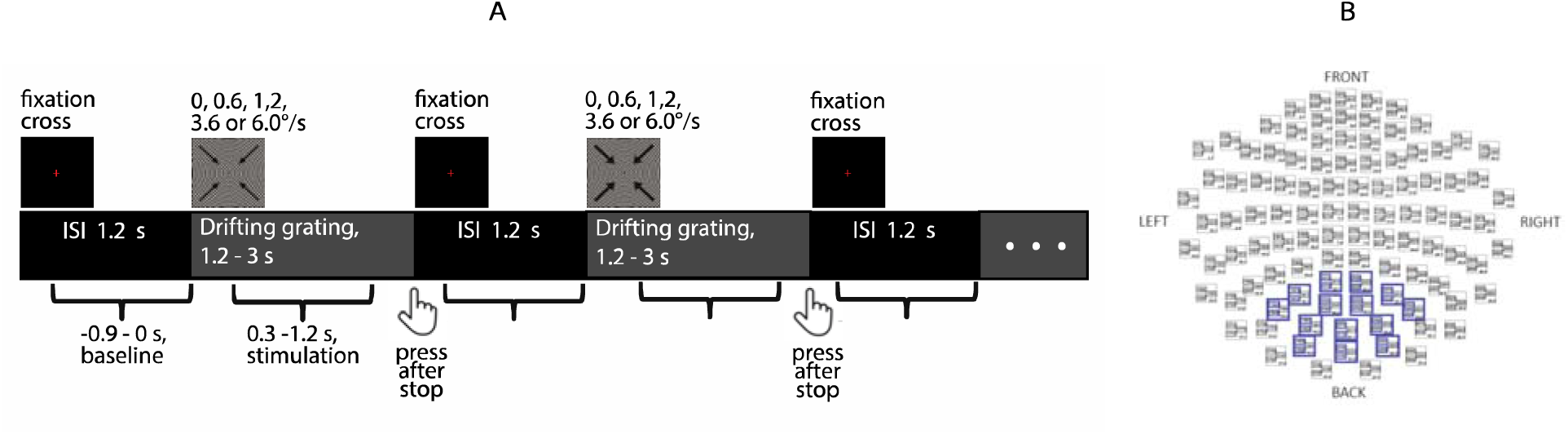
Overview of the visual stimulation paradigm (A) and the arrangement of sensors, among which the ‘maximal gamma sensors’ were selected (B). Curly brackets mark 0.9 seconds intervals included in the time-frequency analysis.

Subjects were offered a break in the middle of the session. For all subjects except one participant with VSS, data were available for both parts of the session: before and after the break (450 trials in total). For one VSS participant, only 200 trials before the first break were available due to a technical error. There was no significant difference between the groups in the duration of the first session (Mann-Whitney U Test, Z = 0.97, Median_VSS_ = 254 trials, Median_Control_ = 258 trials, p = 0.33) or in the break length (Mann-Whitney U Test, Z = 0.64, Median_VSS_ = 41 s, Median_Control_ = 38 s, p = 0.33). In all participants, uninterrupted data were available for 1-137 trials.

### 2.3. Recording of electrophysiological data

The MEG data were recorded at the Moscow Center for Neurocognitive Research (MEG-center, Moscow State University of Psychology and Education) using an Elekta VectorView Neuromag 306-channel MEG detector array (Helsinki, Finland) consisting of 204 planar gradiometers and 102 magnetometers. The MEG signal was acquired with 0.1 Hz high-pass and 330 Hz low-pass built-in filters and sampled at 1000 Hz. Four electrooculogram (EOG) electrodes placed at the outer canthi of the eyes and above and below the left eye were used to record horizontal and vertical eye movements. To monitor heartbeats, electrocardiogram (ECG) electrodes were placed in the area of the sternum and the midaxillary line (ECG lead V6). The subjects’ head position during MEG recording was continuously monitored. Two-way audio and video communication was used to monitor and interact with the subjects in the MEG chamber.

### 2.4. MEG data pre-processing

The raw MEG signal was subsequently processed using MaxFilter software (version 2.2) to minimize interference from external artifact sources through the temporal signal-space separation (tSSS) method^53^ and to provide motion correction. Further data preprocessing steps were performed with the MNE-python toolbox (v.1.7.1). The denoised raw data were notch-filtered at 50:50:250 Hz. Independent Component Analysis (ICA) was then applied for biological artifact correction by detecting and removing independent components (ICs) correlated with heart activity and eye movements. The two groups did not differ in the mean number of ICs removed (N_VSS_ = 2.15; N_control_ = 2.29; Mann-Whitney U Test, Z = 0.41, p = 0.67). The MEG data were epoched from −1 s to 1.2 s relative to stimulus onset. Epochs affected by bursts of myogenic activity or high-amplitude artifacts were identified first through automatic detection of high-amplitude signals (for magnetometers > 4e-12T; for gradiometers > 4000e-13T) and then through visual inspection, and subsequently excluded from the analysis. The average number of artifact-free epochs for the VSS/control group was 83.5/85.6, 83.3/84.2, 82.7/85.6, 83.7/84.9, and 82.9/84.7 for drift rate conditions 0°/s, 0.6°/s, 1.2°/s, 3.6°/s, and 6.0°/s, and did not differ for any of these conditions (all p’s>0.07). Only data from planar gradiometers were used for subsequent analysis.

### 2.5. MEG data analysis

#### 2.5.1. Gamma oscillations: mean power and frequency of gamma response (GR)

Gamma response (GR) power was analyzed within a stimulation window spanning from 0.3 to 1.2 seconds post-stimulus onset. The interval from −0.9 to 0 seconds was used as a baseline. To isolate stimulus-related induced activity, the averaged evoked responses were subtracted from single-trial data according to stimulus type. Time-frequency analysis was performed using the multitaper method with a bandwidth of 10 Hz, a time step of 2 ms, and a frequency resolution of 2.5 Hz. We estimated normalized response power as (stimulation−baseline)/baseline*100%. We then averaged spectra across sensors (1 - 4 sensors) where the average normalized power in the 35-80 Hz range exceeded 80% of that in the ‘maximal sensor’. To minimize the contribution of noise, these sensors were selected from the posterior set of gradiometers (Fig. 1B), where maximal visual GR power is typically expected^54^. The two groups of participants did not differ in the number of sensors selected for averaging (N_VSS_ = 2.69; N_control_ = 2.41; Mann-Whitney U Test, p=0.83).

To estimate the subject’s mean weighted GR power and frequency we used the approach employed in our previous studies (e.g.^28^). Spectra were averaged for all artifact-free epochs, separately for drift rate conditions. The GR power was then calculated as the average of those spectrum values that exceeded 2/3 of the peak value in the frequency range 35–100 Hz. The GR frequency was estimated as the center of gravity of the spectrum values used to calculate the GR power.

#### 2.5.2. Gamma oscillations: single trial GR power and frequency

For the single-trial analysis, GR power and frequency were estimated separately for each trial. For this purpose, the discrete frequency bin closest to the individual’s weighted peak frequency was determined and the power was averaged over a range of ±10 Hz relative to this bin. Single trial GR power can occasionally take negative values because it is calculated as a change relative to the baseline. Therefore, the minimum value was subtracted from the power spectrum before calculating the weighted frequency over the same frequency range of ±10 Hz.

#### 2.5.3. Event-related fields (ERF)

To analyze the time courses of ERF components amplitudes, we first identified the time windows during which individual ERF components were observed. The signal was low-pass filtered at 40 Hz and averaged over all trials, separately for each subject, after which the root mean square (RMS) of the signal across all subject’s gradiometers was calculated. RMS values were averaged over all participants so that VSS and control groups contributed equally to the mean. The mean RMS signal had two distinct peaks with maxima at 84 and 179 ms (M80 and M180) (Figure 5A). Based on the latencies of these peaks, we used time windows of 60-100 ms and 150-240 ms to estimate individual peak latencies for M80 and M180. To this end, we found the gradiometer with the maximal absolute peak amplitude in corresponding time windows. In the maximal peak was negative, the signal was flipped. Single-trial amplitudes were calculated as averages over time windows defined as M80_latency_±15 ms and M180_latency_±30 ms. We used averaging in these time windows rather than detecting peaks in the single-trial data to ensure that component amplitudes were not systematically influenced by changes in the ongoing alpha oscillations, whose power increases with increasing time-on-task^55^.

The peak latencies of the M80 and M180 components averaged across conditions did not differ between subjects with VSS and controls (M80: Mean_VSS_ = 84 ± 8.8 ms, Mean_control_ = 84 ± 8.6 ms, t(52) = 0.02, p = 0.98; M180: Mean_VSS_ = 180 ± 13.8 ms, Mean_control_ = 179 ± 16.2 ms, t(52) = 0.15, p = 0.88). Since we were primarily interested in repetition-related changes in component amplitudes, and the amplitudes were averaged over fairly wide time windows, we did not account for possible differences in component latencies between conditions in this study.

### 2.6. Heart rate variability (HRV)

HRV was assessed using the Systole v0.2.4 package^56^. Two periods of data were analyzed: (1) a 5-minute rest period with eyes open and (2) the first block of the experiment, before the break. ECG data were processed to identify R-peaks using a moving average algorithm. Artifacts such as missed/additional peaks or ectopic beats were corrected using the ‘correct_rr’ function. ECG parameters assessed included heart rate in beats per minute (BPM), as well as basic HRV parameters in the time and frequency domains. The latter included mean standard deviation of R-R intervals (SDNN), mean square of serial differences (RMSSD), percent of successive differences larger than 50 ms (pNN50), high-frequency (0.15-0.40 Hz) power of the R-R interval spectrum (HF), and HF power in normalized units: HFnu = HF/(HF + LF), where LF is the power in the low-frequency range (0.04 to 0.15 Hz). Although all of these HRV measures reflect parasympathetic activity, only HFnu accounts for individual and contextual differences in overall variability and allows us to assess the relative contribution of parasympathetic activity to autonomic balance, regardless of the absolute BMP of HRV levels^49,50^.

### 2.7. Statistical analysis

A mixed ANOVA with Group and Condition factors (drift velocities of 0, 0.6, 1.2, 3.6, and 6.0°/s) was used to analyze group differences in mean GR power and frequency. GR power values were pre-logarithmized to normalize the distributions. If necessary, a Greenhouse-Geisser (G-G) correction was applied to correct for violation of the sphericity assumption. The same ANOVA was used to compare group differences in reaction time. The Wilcoxon rank-sum test was used to compare group differences in error rates.

To analyze the amplitudes of single-trial GR or ERF components, we used the first 137 epochs, which were available for all participants without interruption. We first excluded trials with extremely high or low gamma power or ERF amplitude for each subject based on the interquartile range approach^57^, with a multiplication coefficient of 2. To estimate general patterns of repetition-related changes, single-trial GR power or ERF amplitude values were z-transformed separately for each subject and stimulus type and then averaged across all subjects and stimulus types according to the trial order number. The resulting average was then visually inspected for trends.

Because the averaged z-transformed GR power exhibited the clear biphasic changes described previously for the repetition of static stimuli^41^, we followed the strategy used by the previous authors and fitted the time course of GR power as the sum of exponential and linear processes over the ordinal stimulus number: y ∼ A * exp(-trialN/tau) + B * trialN + C, where tau is the time constant of early gamma decrease and trialN is the trial order number. To better determine the inflection point, we also fitted a broken line model to the same data using the ‘segmented” package in R. We further analyzed only repetition-related changes after the inflection point. The time courses of GR frequency and ERF amplitude were fitted using linear functions. These fits are presented in more detail in the Results section.

Linear mixed model (LMM) analysis, implemented using the ‘lme4’ package in R, was employed to assess factors affecting variations in GR power and ERF amplitude. The models incorporated fixed factors of trial order number (trialN) or the interaction of trialN and group. Random factors (subject and condition) were included with both intercepts and slopes when feasible and appropriate. Model comparison and selection were based on the Akaike information criterion (AIC), Bayesian information criterion (BIC), and Log-Likelihood (logLik) test.

To quantify individual changes in GR power and ERF amplitudes across trials, we employed a subject-specific linear regression model using z-transformed values (z_score ∼ trialN) for each drift rate condition. The resulting coefficients were averaged across conditions per subject, yielding a personalized metric of repetition-related changes in GR power.

For correlation analysis, we used the Spearman method, which is less sensitive to outliers than the Pearson method and does not assume a linear relationship between variables. The Mann-Whitney U Test was used for group comparison of HRV metrics due to their non-Gaussian distribution. Student’s t-test was used for group comparison of normally distributed data.

## 3. Results

### 3.1. Behavioral performance

Reaction time (RT) data were available for 23 of 26 VSS and 26 of 27 control participants with 4 subjects’ data lost due to technical issues. Omission error rates were comparable between groups (VSS median: 1.2%, Control median: 0.96%; Wilcoxon W = 252.5, p = 0.42). However, commission errors were significantly higher in the VSS group compared to controls (Median_VSS_ = 4.1%, Median_Control_ = 2.0%; Wilcoxon W = 390, p = 0.006). This suggests that while both groups showed similar task engagement, VSS participants exhibited greater difficulty in response inhibition, potentially indicating differences in cognitive control mechanisms between the two groups.

Analysis of correct trials using mixed ANOVA revealed a significant effect of Condition on RT (F(4, 188) = 10.33, G-G ε = 0.47, p = 0.0001, η = 0.18), with RT decreasing as drift rate increased. However, neither Group effect (F(2, 47) = 0.28, p = 0.59, η = 0.01) nor Group x Condition interaction (F(4, 188) = 1.53, G-G ε = 0.47, p = 0.20, η = 0.03) reached significance. These results suggest that while drift rate significantly influenced response times, the performance pattern was similar across groups.

### 3.2. No group differences in grand average gamma power, frequency or their modulation by drift rate

Mixed ANOVA analysis of GR power averaged across all trials, with Group and Condition as factors and normalized Age as a covariate, revealed a robust effect of Condition (F(4, 200) = 81.0, G-G ε = 0.43, p < 1e-6, ηp^2^ = 0.62; Fig. 1A). In accordance with previous results^19^, an average GR power increased with the drift rate, peaking at 1.2°/s, before declining as the drift rate exceeded this value. However, neither Group effect (F(1, 50) = 0.68, p = 0.41, ηp^2^ = 0.01) nor Group x Condition interaction (F(4, 200) = 0.69, G-G ε = 0.43, p = 0.6, ηp^2^ = 0.01) reached significance. Notably, individual data examination uncovered substantial inter-individual variability in GR power (Fig. 1B).

For the average GR frequency, mixed ANOVA also showed a highly significant effect of Condition (F(4, 200) = 325.9, G-G ε = 0.37, p < 1e-6, ηp^2^ = 0.86; Fig. 1C), but no significant effects of Group (F(1, 50) = 0.81, p = 0.37, ηp^2^ = 0.02) or Group x Condition interaction (F(4, 200) = 0.32, p = 0.86, ηp^2^ = 0.01). In both groups, mean GR frequency consistently increased with the drift rate, reaching its maximum at 6.0°/s^19^. Gamma frequency significantly decreased with age (F(1, 50) = 5.24, p = 0.026, ηp^2^ = 0.09), aligning with previous findings ^19,58^.

### 3.3. GR power shows a biphasic pattern with stimulus repetition

Analysis of the z-transformed GR power averaged over all participants according to the trial number revealed a distinct biphasic pattern: a rapid initial decrease followed by a near-linear increase. Figure 3 illustrates this pattern over 137 trials, modeled as a combination of exponential decay and linear increase (R²adj = 0.74). The early decrease exhibited a time constant tau = 7.8 repetitions (CI95% = [6.0, 9.6]). Separate analyses for VSS (R²adj_VSS_ = 0.54, tau_VSS_ = 7.3, CI95%_VSS_ = [3.7, 10.7]) and control groups (R²adj_control_ = 0.40, tau_control_ = 7.7, CI95%_control_ = [4.7, 10.7]) demonstrated similar biphasic patterns. A broken-line fit estimated the inflection point at 14.6 trials (CI95% = [12.6, 16.6]). Given our focus on the later repetition effect, potentially linked to Hebbian plasticity^38,41^, we subsequently limited the analysis to trials 15-137.

Previous studies have highlighted the stimulus dependency of repetition-related GR changes^38,41^. Although the visual stimulus in our study had the same shape and contrast across trials, its drift rate varied between trials. The limited amount of data prevented us from testing the effects of the trial number and drift rate repetition within a single model. To determine which factor better explained the time-dependent changes in GR power, we compared two LMMs, each of which accounted for random intercepts and slopes for individual subjects and conditions:

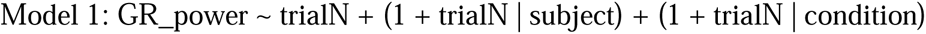

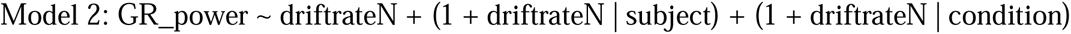

Model 1 examined the effect of overall trial number (15-137), while Model 2 focused on drift-rate-specific repetition number within the same range. Model 1 demonstrated superior fit (AIC = 13611, BIC = 13671, logLik = −6796.3) compared to Model 2 (AIC = 13674, BIC = 13734, logLik = −6827.8). These findings indicate that trial order number was a more effective predictor of z-transformed GR power than the repetitions of specific drift rate in our study design.

We then expanded Model 1 to include a trialN * group interaction:

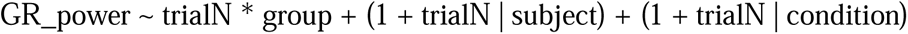

Analysis revealed a large and highly significant trial order number (trialN) effect: t(20.90) = 5.01, p < 0.0001, d = 1.10. There was also significant trialN x group interaction: t(51.15) = 2.26, p = 0.028, d = 0.32 due to steeper GR power increase with trial number in participants with VSS. The effect of the group was non-significant: t(50.98) = 1.59, p = 0.118, d = 0.22, indicating an absence of notable group differences in average GR power across trials 15-137.

Although our study design did not allow us to analyze the effect of drift rate on the stimulus repetition-related increase in GR power, inspection of the median power values in the trials averaged within 40-trial blocks (trials 15 - 55, 56 - 96, 97 - 137) showed that the GR power in the third block exceeded that in the first block for all drift rates and in both groups of participants (Fig. 3D).

To rule out baseline gamma power changes as a driver of group differences in stimulus repetition-related GR power increases, we applied the model:

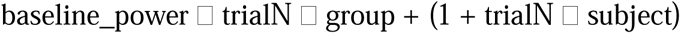

Results revealed no significant effect of group (t(47.00) = 0.30, p = 0.76, Cohen’s d = 0.043) or trialN * group interaction (t(46.37) = 0.80, p = 0.43, Cohen’s d = 0.118). These findings indicate that the observed group differences in repetition-related GR power increases are not attributed to baseline changes.

To quantify individual GR power changes between trials 15-137, we fitted linear regression models [z_GR_power ∼ stimN] for each subject and drift rate condition, then averaged coefficients across conditions. Both groups showed significant non-zero averaged coefficients, VSS: mean = 0.246, SD = 0.160, t(25) = 7.84, p < 1e-6, Cohen’s d = 1.57; Control: mean = 0.168, SD = 0.152, t(26) = 5.72, p < 0.00001, Cohen’s d = 1.12 (Fig. 3C). Mann-Whitney U Test confirmed group differences in regression coefficients (N_VSS = 26, N_Control = 27, Z = 2.14, p = 0.03, r = 0.29, medium effect), indicating stronger repetition-related GR power increases in VSS participants.

To test for a possible link between average GR power and time-dependent GR power changes we calculated the correlation between the GR power averaged over all trials and conditions and the time-dependant regression coefficients descrived above. A significant direct correlation was observed only in VSS participants (Spearman correlation: VSS: N = 26, R = 0.39, p = 0.046), while no significant correlation was found in the control group (N = 27, R = −0.06, p = 0.76).

In summary, both control and VSS participants exhibited a rapid decrease in GR power during the initial trials, followed by a sustained increase over the subsequent 100+ trials. Notably, VSS participants showed a more pronounced increase in GR power, which correlated with their mean GR power—a pattern absent in control participants. Given that the same grating was presented across trials with varying drift rates, we will hereafter refer to the trial-by-trial changes in GR power as ‘repetition-related GR changes’.

### 3.4. Stimulus repetition-related GR power increase is associated with HRV

First, we examined group differences in time-based HRV metrics (RMSSD, SDNN, pNN50), frequency-based HRV metrics (HF, HFnu), and mean heart rate (BPM). No significant group differences were found in HRV metrics during either resting with eyes open or during visual stimulation (Mann-Whitney U Test, all p > 0.22). However, during rest, the VSS group showed a trend towards a higher BPM compared to the control group (N_VSS_ = 26, N_Control_ = 23, Z = 1.74, p = 0.08). We then investigated the relationship between individual GR power regression coefficients and HFnu, a measure that reflects the relative contribution of parasympathetic activity to autonomic balance, independent of heart rate and absolute HRV levels^49,50^.

During rest, GR power regression coefficients showed a significant correlation with HFnu in the overall sample (N = 49, R = 0.32, p = 0.03) and in the control group (N = 23, R = 0.53, p = 0.01). A similar trend was observed in the VSS group (N = 26, R = 0.35, p = 0.08). During visual stimulation, the correlation between GR power regression coefficients and HFnu remained significant in the combined participant sample (N = 53, R = 0.30, p = 0.03) and in the VSS group (N = 26, R = 0.53, p = 0.006), but not in the control group (N = 27, R = −0.12, p = 0.55). Figure 4 illustrates the relationship between GR power regression coefficients and HFnu. Additional correlations between GR power regression coefficients and all HRV measures are detailed in *Supplementary table S1*.

### 3.5. No correlation between repetition-related GR power increase and symptoms severity

There was no correlation between the slope coefficient of repetition-related GR power changes and scores on the Visual Discomfort scale in patients with VSS (N = 24, Spearman R = 0.027). Among the VSS participants, most had a history of migraine, and there were no significant differences in GR power slope coefficients between those with and without migraine (With migraine: N = 21, mean b = 0.24; Without migraine: N = 5, mean b = 0.29; Mann-Whitney U Test: Z = 0.39, p = 0.70). Additionally, VSS subjects with migraine with aura did not differ from those with migraine without aura in this regard (Migraine with aura: N = 7, mean b = 0.26; Migraine without aura: N = 24, mean b = 0.22; Mann-Whitney U Test: Z = 0.79, p = 0.43).

### 3.6. Repetition-related increase in GR frequency does not differentiate between VSS and control groups

For the combined sample, a model accounting for random intercepts and slopes for individual subjects and a random intercept for stimulus type [GR_frequency ∼ trialN + (1 + trialN | subject) + (1 | condition)], provided a better fit than the more complex model, which included random intercepts and slopes for both subjects and stimulus type [GR_frequency ∼ trialN + (1 + trialN | subject) + (1 + trialN | condition)]. The fit statistics were as follows: simple model AIC = 33776, BIC = 33823, logLik = −16881; complex model AIC = 33780, BIC = 33839, logLik = −16880. Therefore, for simplicity, the slope for a condition factor was not included in the model. With this LMM, the effect of stimulus repetition within trials 15-137 was significant (t(49.34) = 2.51, p = 0.015, Cohen’s d = 0.35) due to an increase in GR frequency with time.

The inclusion of the group factor in this model [ GR_frequency ∼ trialN * group + (1 + trialN | subject) + (1 | condition) ] revealed no significant effects for either the group or the trialN * group interaction. Specifically, the results showed no significant group effect (t(50.0) = 0.52, p = 0.60, Cohen’s d = 0.06) and no significant interaction between trialN and group (t(48.6) = 1.21, p = 0.23, Cohen’s d = 0.17).

In conclusion, the peak frequency of the oscillatory gamma response increased with the number of stimulus repetitions, but this trend did not differ significantly between the VSS and control participants.

### 3.7. Habituation of evoked responses does not differentiate between VSS and control groups

First, we examined potential group differences in RMS values of the averaged signal within the M80 and M180 time windows (M80 latency ± 15 ms and M180 latency ± 30 ms) using a mixed ANOVA with factors of Group and Condition. No significant effects of Group or Group × Condition interaction were found for either component (all p > 0.3). However, the effect of Condition was significant for M80 (F(4, 204) = 3.9, G-G ε = 0.93, p = 0.004, η = 0.07), which showed a decrease with increasing drift rate, and for M180 (F(4, 204) = 13.8, G-G ε = 0.49, p = 0.000006, η = 0.21), which exhibited an increase at higher drift rates (Figure 5 B, C).

To analyze the temporal dynamics of M80 and M180 responses across the entire sample of participants, we employed a method similar to that used for GR parameters. First, amplitude values were z-transformed separately for each subject and condition, then averaged across all conditions and participants for visual inspection. The z-scaled M80 amplitudes showed a clear decreasing trend with increasing number of trials, reaching a plateau around the 60th sample (Fig. 5D). To identify the inflection point of this curve, we fitted a split line to the data, excluding the first trial due to its anomalously low M80 z-scored amplitude (Figure 5D). This analysis pinpointed the inflection point at the 57th trial. Based on this finding, we applied a LMM to the single-trial M80 amplitudes for trials 2 - 57. This model accounted for slope and intercept for individual subjects and intercept for condition: [M80ampl ∼ trialN + (1 + trialN | subject) + (1 | condition)]. The results revealed a significant effect of stimulus repetition (t(52.01) = 3.43, p = 0.001, Cohen’s d = 0.48), indicating robust habituation. To investigate potential group differences, we expanded the model to include a fixed effect of group: [ M80ampl ∼ trialN * group + (1 + trialN | subject) + (1 | condition) ]. There was a trend towards less pronounced M80 amplitude habituation in the VSS group (t(51.17) = 1.70, p = 0.096, Cohen’s d = 0.24). This subtle difference is illustrated in Figure 5E using z-scored data.

The z-scored M180 amplitudes exhibited a nearly linear decrease with increasing trial order number, with a notable exception for the first four trials, which deviated from this habituation pattern (Fig. 5F). To account for this initial irregularity, we excluded these four trials from the subsequent analysis. We then applied the LMM to the single-trial M180 amplitudes for trials 5- 137, using the following model: [ M180ampl ∼ trialN * group + (1 + trialN | subject) + (1 | condition)]. This analysis revealed a significant effect of trial order number (t(51.89) = 3.36, p = 0.001, Cohen’s d = 0.47), indicating a robust habituation effect across the combined sample of participants. The interaction between a trial order number and group was not significant (t(50.95) = 1.03, p = 0.31, Cohen’s d = 0.14). These results, illustrated in Figure 5G using z-scored data, suggest that while there is a clear habituation effect in M180 amplitude across all participants, this effect does not significantly differ between the VSS and control groups.

In conclusion, our analysis revealed that neither M80 nor M180 amplitude habituation significantly differentiated between VSS and control groups. However, a subtle trend towards reduced M80 amplitude habituation was observed in VSS participants, suggesting a potential difference in neural adaptation processes.

### 3.8. The relationship between repetition-related changes in GR power and EFR components

To investigate a potential link between the habituation of evoked components and stimulus repetition-related changes in GR power, we first quantified these changes in M80 and M180 using the same approach as for GR power. Specifically, we fitted linear regressions to the z-transformed amplitude values for each drift rate condition [M80amplitude ∼ trialN, M180amplitude ∼ trialN] and then averaged the linear regression coefficients across conditions for each subject.

For both groups, coefficients were significantly different from zero for the M80 component (VSS: mean = −0.075, SD = 0.156, t(25) = 2.46, p = 0.02, Cohen’s d = 0.48; Control: mean = - 0.170, SD = 0.200, t(26) = 4.41, p = 0.0002, Cohen’s d = 0.85) and M180 component (VSS: mean = −0.075, SD = 0.168, t(25) = 2.27, p = 0.03, Cohen’s d = 0.45; Control: mean = −0.116, SD = 0.190, t(26) = 3.17, p = 0.004, Cohen’s d = 0.61).

In the control group, the correlations between evoked response regression coefficients and GR power regression coefficients were not significant (N = 27; M80: R = −0.02, M180: R = 0.07). The VSS group showed a trend towards a positive correlation between M80 and gamma power regression coefficients (N = 26; R = 0.35, p = 0.08), suggesting a potential link between weaker M80 habituation and stronger gamma increases in VSS participants. The M180 regression coefficient did not correlate with the gamma slope in the VSS (N = 26; R = −0.02, n.s.) or control (N = 27; R = 0.07, n.s.) groups.

## 4. Discussion

In this MEG study, we investigated potential abnormalities in visual oscillatory gamma response (GR) parameters in patients with Visual Snow Syndrome (VSS) that reflect the excitation-inhibition balance and neuroplasticity in the early visual cortex. Our results indicated no significant differences between VSS and control groups in average GR power or frequency, nor in their modulation by varying visual stimulation intensity, suggesting an intact excitation-inhibition ratio in VSS. Both groups exhibited biphasic changes in GR power with stimulus repetition: an initial rapid decrease over the first few trials, followed by a gradual increase across approximately 100 subsequent repetitions. Notably, VSS patients demonstrated a significantly amplified GR power increase during the second phase. These findings imply that enhanced neuroplasticity in the early visual cortex may play an important role in the pathophysiology of VSS, offering new insights into the underlying mechanisms of this condition.

The biphasic changes in single-trial visual GR power with stimulus repetition observed in our study (Figure 3A) closely resemble those recently described for the first time in MEG of healthy human subjects^41^. Stauch and colleagues demonstrated that repetitive presentation of an oblique grating initially led to a rapid decrease in visual GR power (time constant: 3.5 repetitions), followed by an approximately linear increase over the subsequent ∼100 repetitions. Importantly, these repetition-related power changes were stimulus-specific: altering the grating orientation disrupted the pattern, confirming their genuine link to neuronal plasticity rather than confounding factors (e.g. fatigue) that might influence neural responsiveness over time. The stimulus specificity, coupled with the persistence of the effect over several minutes, provides strong evidence for the involvement of experience-dependent plasticity mechanisms in shaping gamma oscillatory responses to repeated stimuli.

In our study, the annular grating pattern remained constant, while its drift rate varied randomly across trials. This additional variability may explain the slower initial habituation of GR power observed in our study in both groups, with a time constant of 7 – 8 repetitions. The slope of the subsequent repetition-related increase in GR power was positive for nearly all participants (Figure 3C), indicating a reliable repetition-related gamma increase. Consistently with previous findings, we also observed a repetition-related increase in GR frequency.

These repetition-related GR changes are thought to reflect Hebbian plasticity, characterized by strengthened synaptic connections among the ‘most responsive’ neurons and weakened connections among the ‘less responsive’ ones^38,41^. Studies in monkeys using peripheral grating presentations demonstrated that the repetition-related increase in GR power was accompanied by an initial decrease and a subsequent stabilization in neuronal spiking activity^38,59^. This pattern suggests that the observed GR power increase reflects enhanced neural synchronization rather than elevated neural excitation. Conversely, Galuske et al.^37^ demonstrated in cats that an increase in gamma power induced by repeated exposure to a full-field grating with a specific orientation did not alter the firing rates of neurons tuned to that orientation. However, it did increase the activity of neurons tuned to similar orientations within a 30° range.

Importantly, both described neuronal mechanisms suggest that stimulus repetition enhances the involvement of V1 neuronal populations in synchronous gamma oscillations, thereby strengthening their collective neural impact on postsynaptic target neurons in downstream cortical areas^41,60–63^.

From this perspective, the enhanced repetition-related gamma synchronization in V1 of patients with VSS may contribute to overstimulation of their downstream secondary and associative visual areas. Metabolic abnormalities in these regions, both during rest and visual stimulation, are consistently reported in this neurological disorder^9,11,13^. Our results suggest that the observed hypermetabolism in higher-tier visual areas in VSS may thus be a consequence of the amplified input from V1, reflecting a cascade effect of altered neural processing throughout the visual hierarchy.

Regardless of precise mechanisms, the enhanced repetition-related changes in gamma response power observed in VSS participants likely reflect heightened Hebbian plasticity associated with activity-dependent modifications in neural representations of frequently encountered visual stimuli. Notably, animal studies have underscored the causal role of the maladaptive neural plasticity in the generation of tinnitus^64^, an auditory misperception disorder often considered analogous to VSS (e.g.,^30^). Given our findings, the high comorbidity of tinnitus in VSS patients suggest a shared pathophysiological mechanism involving altered neural plasticity.

The association between repetition-related gamma enhancement and the cardiac-derived index of autonomic balance provides additional indirect support for the ‘plasticity interpretation’ of our gamma findings. Our study revealed that a greater shift towards parasympathetic autonomic activity correlated with a steeper repetition-related increase in GR power (Figure 4). States characterized by parasympathetic dominance appear to facilitate neuroplasticity and cortical remapping of representations of repeated stimuli. Indeed, numerous human and animal studies demonstrate that chronic or acute parasympathetic activation, such as through a vagus nerve stimulation, enhances stimulus-specific plasticity across sensory modalities by activating multiple neuromodulatory systems^44,65–67^.

Acetylcholine – the main neurotransmitter in the parasympathetic autonomic system – plays an important role in both neuroplasticity^43^ and gamma oscillations^68^. In cats, pairing repetitive visual stimulation with stimulation of the cholinergic nuclei in the mesencephalic reticular formation— mimicking arousal—led to enhanced gamma oscillations and long-term remapping of V1 cortical neurons^37^. Similarly, stimulation of the cholinergic nucleus basalis has been found to support activity-dependent plasticity in the primary auditory cortex during repeated auditory stimuli^69^. These studies underscore the dependency of neuroplasticity in the early sensory cortical areas on functional states, such as cholinergic arousal and attention.

The observed correlation between the shift toward parasympathetic activity, as indicated by the HRV index HFnu, and repetition-related changes in GR power aligns with the role of functional states associated with parasympathetic activation (e.g., selective attention^70^) in promoting neuronal plasticity^71^. However, our study found no differences in HRV parameters between participants with VSS and controls, suggesting that the observed group differences in the repetition-related GR power slope are more likely trait-like in nature rather than driven by state-related variations in autonomic activity.

Our results suggest that aberrant neuroplasticity may contribute to VSS pathophysiology; yet we found no significant correlation between an increase in repetition-related gamma synchronization (a marker of enhanced plasticity) and the severity of visual discomfort or migraine comorbidity in patients with VSS. This indicates that the observed neural changes in the early visual cortex, a primary generator of visual gamma oscillations^17,18^, may not directly relate to other neurological alterations common in VSS patients. Similar dissociations exist in tinnitus, where self-reported distress correlates more with the activity of the stress-related brain regions and their connectivity to the auditory cortex rather than with sensory cortex activation^72–74^. Further research is necessary to determine whether the enhanced plasticity in early visual cortical areas observed in VSS patients predisposes them to the disorder’s core symptom—phantom visual ‘snow’ sensations.

Our findings regarding alterations in neuroplasticity in VSS may have valuable clinical implications for therapeutic interventions for this disorder. They suggest that targeting neuroplasticity through neuropharmacological, instrumental, or psychotherapeutic approaches may be effective in alleviating both visual and psychological symptoms associated with this condition. Although evidence for the efficacy of such treatments in VSS is still limited, there are some promising examples. For instance, up to 20% of VSS patients report benefits from lamotrigine^75,76^, an antiepileptic drug that not only reduces neuronal excitability by inhibiting glutamate release but also attenuates long-term potentiation (LTP) and long-term depression (LTD)-like synaptic plasticity^77^. Additionally, Mindfulness-Based Cognitive Therapy (MBCT), which has been shown to induce functional and structural brain changes indicative of neuroplasticity^78^, has demonstrated improvements in visual network functional connectivity and behavioral outcomes in VSS patients^79^. Furthermore, transcranial magnetic stimulation (TMS), which can modulate synaptic connections in targeted brain regions and induce plastic changes, has emerged as a promising therapeutic tool for VSS^80^. The neurofunctional indices of neuroplasticity identified in our study could facilitate the selection of patient populations most likely to benefit from these targeted interventions, potentially enhancing treatment efficacy for VSS.

Despite the observed group differences in the repetition-related GR changes, we found no significant differences in the average power or frequency of visually induced GR to static or moving gratings, nor in the modulation of GR parameters by grating drift rate (Figure 2). Given the proposed relationship between parameters of gamma oscillations and the E-I ratio in the visual cortex, these results indicate that any increased neuronal excitability in VSS was not substantial enough to disrupt the characteristics of non-invasively recorded gamma oscillations.

**Figure 2.**
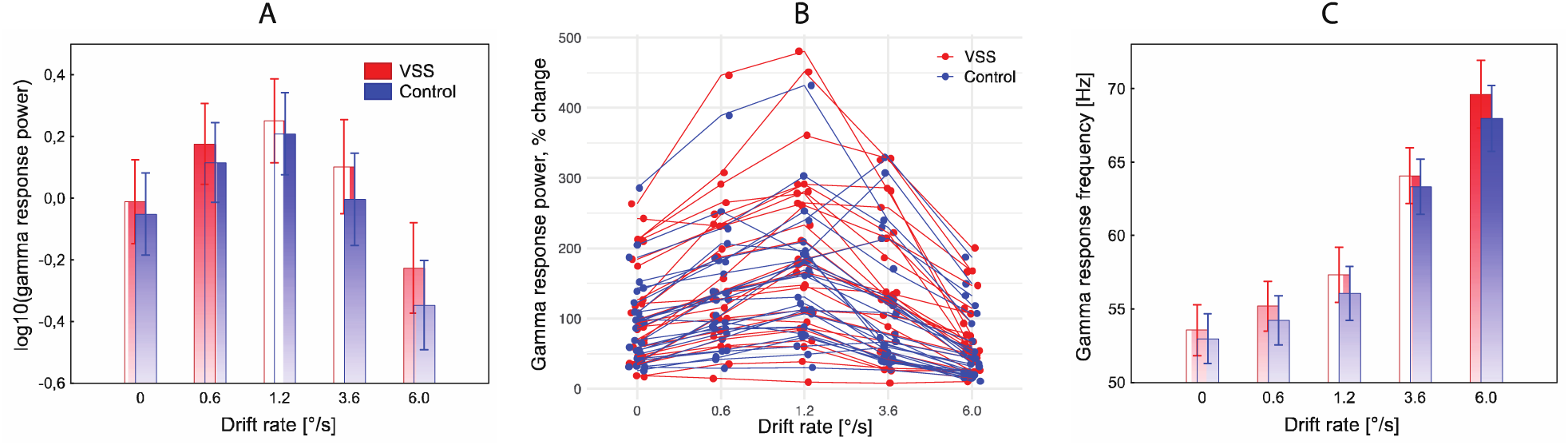
Changes in the spectral parameters of visual gamma response (GR) as a function of grating drift rate in patients with VSS and control participants. **A**. Log-transformed average GR power in the two groups. **B.** Interindividual variability of GR power. **C.** Average GR frequency in the two groups. Vertical bars in (A) and (C) denote 0.95% confidence intervals.

Notably, the only other MEG study that estimated visual gamma oscillations in VSS^14^ reported in patients with this syndrome heightened visually induced gamma oscillations compared to healthy control participants. The authors interpreted this finding as indicative of increased excitability of V1. This discrepancy may reflect the heterogeneity of VSS, where elevated excitability in the primary visual cortex might only occur in a subset of the patients’ population, potentially leading to sampling bias. Interestingly, in our VSS sample, but not in control participants, a greater magnitude of induced gamma synchronization across all drift rates was linked to a stronger repetition-related gamma enhancement. This finding suggests that putatively elevated neural excitability in some individuals with VSS may amplify their abnormal activity-dependent cortical plasticity, aligning with evidence that elevated excitability can modulate and enhance plastic changes in neural assemblies^81^.

A more critical point of debate, surpassing the discrepancy in findings discussed earlier, is whether absolute GR power recorded via MEG reliably reflects neuronal excitability. While some limitations were addressed in the introduction, this issue is particularly significant for interpreting GR findings in clinical populations. From a functional perspective, increases in GR power may signify enhanced neural synchronization rather than an actual rise in neuronal firing rates^59^, which is the definitive indicator of neuronal excitability. Moreover, relying on GR power measured in a single experimental condition, as an indicator of the excitability of an individual’s visual cortex is particularly problematic. GR power is highly sensitive to stimulation parameters^82^, and this sensitivity can vary considerably between subjects and clinical conditions, complicating direct comparisons. Within a group, for instance, a subject may exhibit high GR power in one stimulation condition but low GR power in another. In a recent study, for example, we found no significant correlation between GR power induced by static gratings and that induced by gratings drifting at high speeds^82^.

In this context, assessing the *modulation* of GR parameters by excitatory drive offers a more reliable approach for estimating neural excitability and E-I balance in the early visual cortex. A detailed discussion of this issue is available in our previous studies (e.g.^29^); here, we provide a brief summary. A gradual increase in excitatory drive to V1 leads to a rapid rise in excitation and a slower, concurrent increase in inhibition^83^. According to computational models, at high excitatory drive, when the inhibitory population of neurons is recruited faster than the excitatory population, the power of gamma oscillations begins to decrease^26,27^. Consequently, the degree of GR power reduction when excitatory drive increases beyond the ‘optimal’ point for gamma generation may serve as a marker of neural inhibition efficiency in controlling the increasing excitation within V1 circuitry.

We have previously observed that the modulation of GR power by varying drift rates of a grating (a proxy for excitatory drive) is altered in conditions associated with E-I imbalance^28,29^. For instance, children with autism—characterized by inefficient inhibition in the primary visual cortex^84^—exhibited reduced GR suppression at higher drift rates, indicating an elevated E-I ratio^29^. In this context, the absence of abnormalities in GR power modulation among patients with VSS (Figure 2 A,B) suggests a preserved E-I ratio in their early visual cortex.

Beyond GR power, GR frequency and its modulation by excitatory drive were also found to be normal in participants with VSS (Figure 2C). Animal and computational studies consistently associate the modulation of gamma oscillation frequency (e.g., by stimulus contrast, velocity, or size) with the functionality of fast-spiking PV+ interneurons^21,85,86^. Previously, we documented attenuated GR frequency modulation in children with autism^29^, consistent with findings from animal models indicating PV+ interneuron dysfunction in the primary visual cortex as a characteristic feature of this condition^84^. In contrast, the normal GR frequency and its modulation observed in VSS suggest that the functioning of this class of interneurons involved in gamma generation is preserved.

Taken together, our findings of relatively normal GR frequency and power modulation with increasing input drive in individuals with VSS suggest the absence of a significant inhibitory deficit or notable alterations in the E-I balance within the primary visual cortex. The excitatory-inhibitory recurrent interactions responsible for generating and modulating gamma oscillations in V1 remain largely intact in VSS patients. Even if excitability is elevated, the E-I balance in the early visual cortex in individuals with VSS seems to remain well-compensated, likely due to the preserved function of inhibitory neurons that effectively counteract increased excitation. In this context, the heightened activity-dependent plasticity observed in VSS patients is unlikely to stem from impaired inhibition or an altered excitation-inhibition balance in the early visual cortex. This conclusion is supported by the coexistence of atypical repetition-related gamma power changes with normal responsiveness of gamma oscillations to excitatory drive in this clinical population.

Another question arising from our results is the comparative sensitivity of two neurophysiological indices—facilitation of visually induced gamma oscillations and habituation of phase-locked evoked responses (ERF/ERP)—to potential neuroplasticity abnormalities in the early visual cortex in VSS. In our study, repeated exposure to the stimulus affected both the induced gamma oscillations (Figure 3) and the magnitude of visual evoked responses (Figure 5). ERP/ERF habituation, defined as a decrease in component amplitude upon repetition, is thought to reflect the ‘fine-tuning’ of neurons, optimizing responses to repetitive stimuli and reducing cumulative cortical activation^87^. Previous studies have frequently reported a lack of visual ERP habituation in migraine, a condition often comorbid with HRV, during the interictal period^9^, but see^40^. Although similar trends have been reported in a few VSS studies^6,15,16^, limited results remain inconsistent^88^.

**Figure 3.**
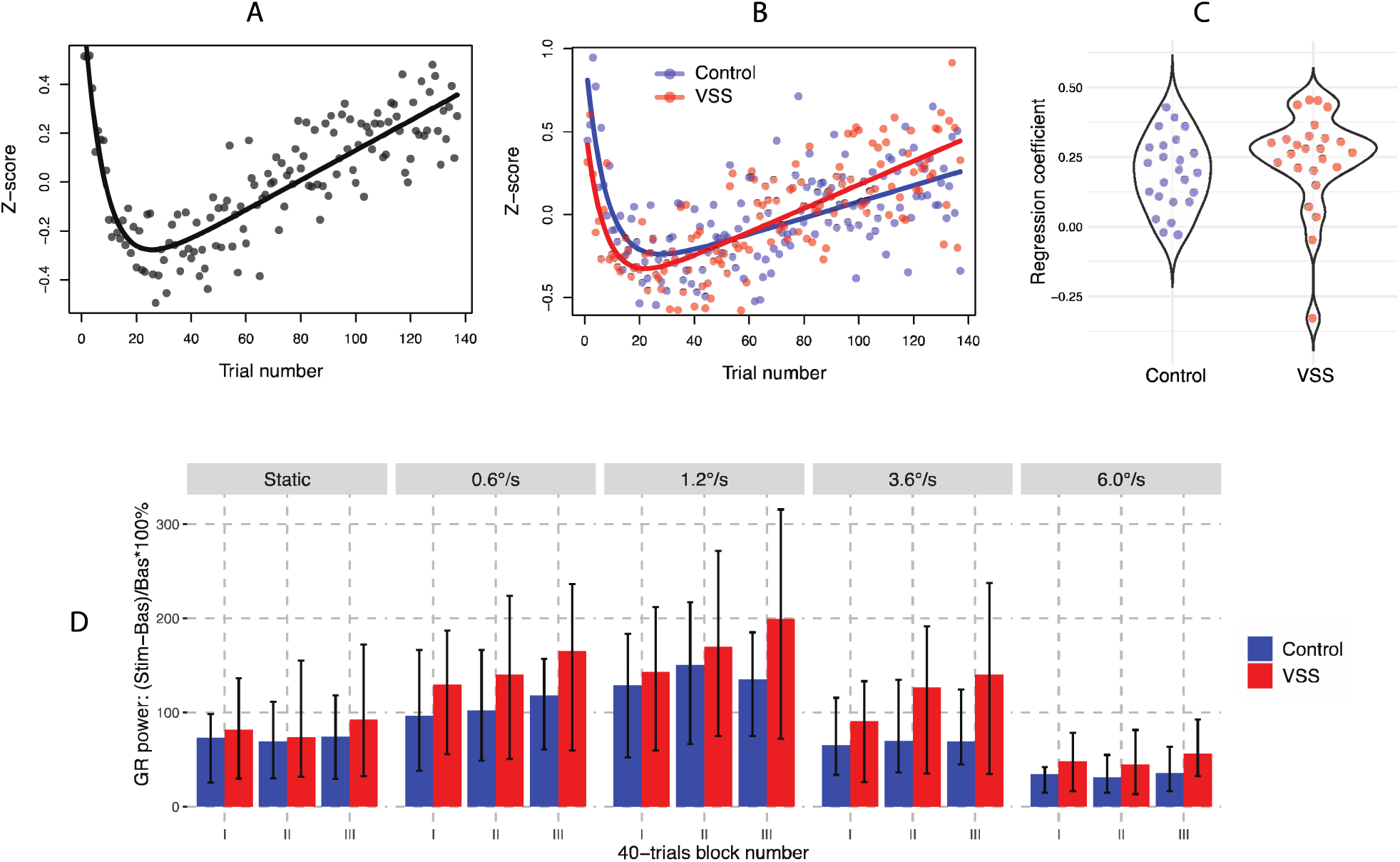
Repetition-related changes of gamma response (GR) power. **A.** Time course of z- transformed GR power in the combined group of participants. **B.** Time courses of z-transformed GR power in VSS and control groups. **C.** Violin plots of individual regression coefficients for 15 −137 trials, averaged across drift rates in control and VSS groups. **D.** Median values of GR power in three blocks of trials (I: 15-55, II: 56-96, III: 97-137) plotted separately for experimental groups and drift rate conditions. Whiskers mark 75% percentile-based confidence intervals.

**Figure 4.**
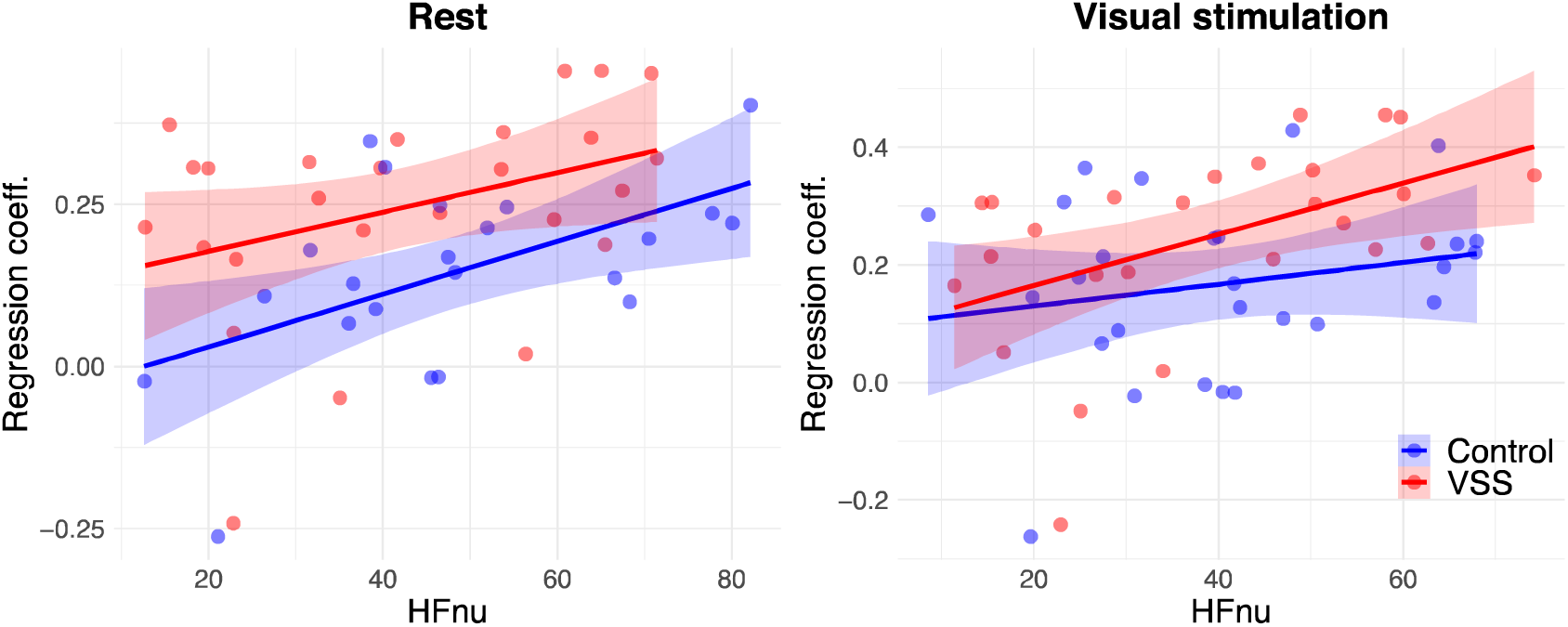
The relationship between repetition-related changes in gamma response power in VSS and Controls, quantified via the mean regression coefficient, and the high-frequency normalized heart rate variability (HFnu) index of autonomic balance.

**Figure 5.**
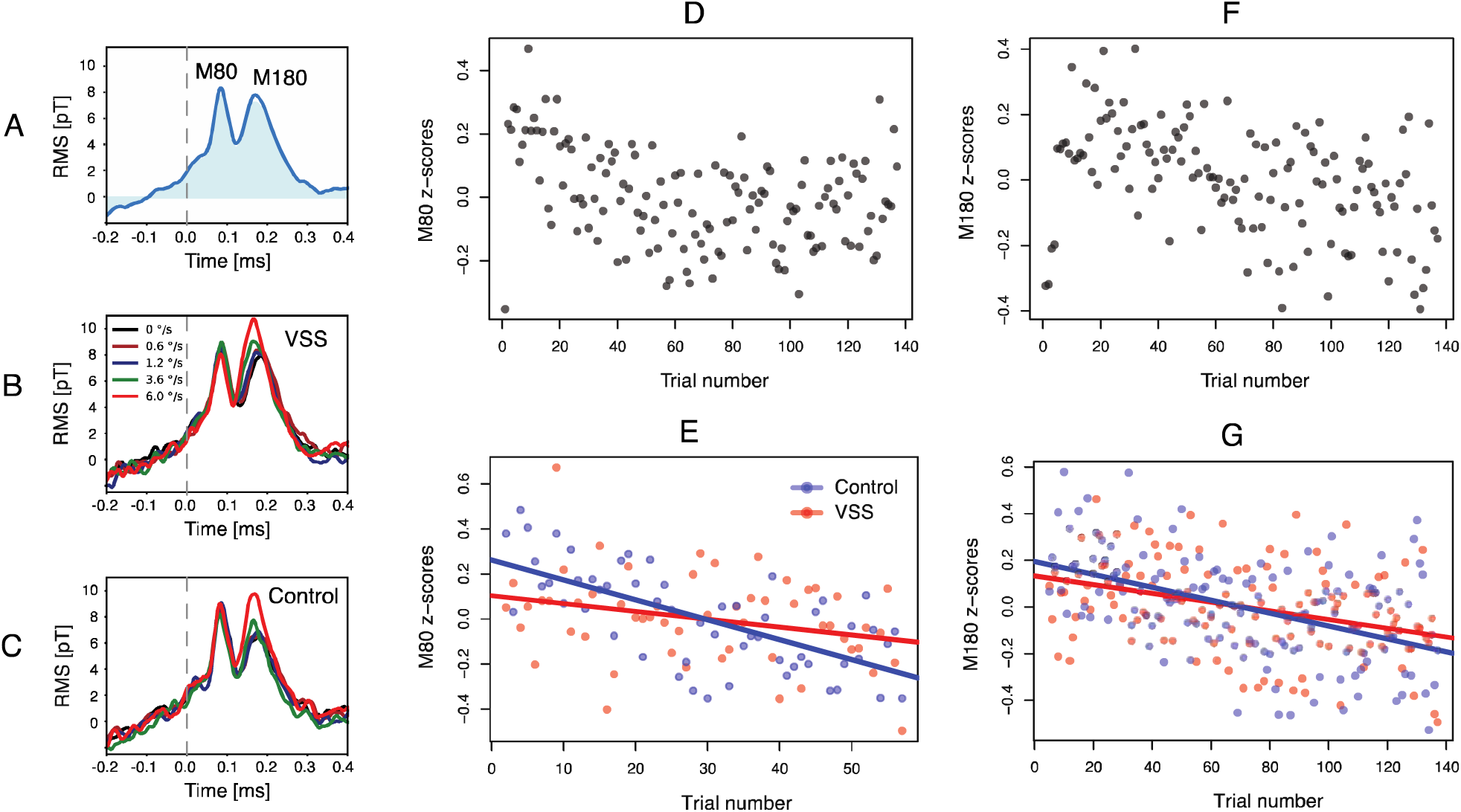
Changes in amplitude of event-related fields (ERF) associated with stimulus repetition. **A.** Root mean squared (RMS) signal averaged over all gradiometers, participants and drift rate conditions. **B, C.** RMS signals averaged separately for drift rate conditions, for VSS and control participants, respectively. **D.** Time course of z-scored M80 amplitude across 1 - 137 trials in the combined sample of participants. **E.** Time courses of z-scored M80 amplitude in VSS and control participants across 2 - 57 trials. **F.** Time course of z-scored M180 amplitude across 1 - 137 trials in the combined sample of participants. **G.** Time courses of z-scored M180 amplitude in VSS and control participants across 5-137 trials.

In the present study, we observed reliable habituation of visual ERF components (M80 and M180) in both control participants and those with VSS (Figure 5). This result resembles the findings of Stauch et al.^41^, who presented healthy participants with static gratings for 0.3 - 2 seconds with 1-second intervals and described a trial-dependent decrease in the magnitude of the ‘early’ and ‘late’ ERF components induced by the gratings (55 - 70 and 90 - 180 ms, respectively, in their study). ERF activity around ∼70 - 80 ms is likely, at least partially, generated in the early visual cortex^89^, which allows for comparison of its repetition-related changes with those observed in GR power. In our study, the magnitude of the M80 peak gradually decreased during the first ∼60 stimulus repetitions and then reached a plateau (Figure 5D). This time course contrasts with the dynamics of GR power, which decreased rapidly during the first few presentations and then gradually increased over at least 100 trials (Figure 3A). VSS patients exhibit a trend toward attenuated M80 habituation, along with a tendency for correlation between reduced M80 habituation and increased GR facilitation during the later phase of stimulus repetitions. Although both M80 findings did not reach statistical significance, we report them to emphasize the need for future studies with larger sample sizes and more refined experimental designs to clarify the relationship between ERF habituation and gamma amplification in VSS. However, given the different trial-wise dynamics of these electrophysiological measures, they are likely driven by separate, co-occurring repetition-related mechanisms within the early visual cortex. Notably, only gamma synchronization exhibited atypical repetition-related patterns in VSS in the present study.

Our study has several limitations. The experimental design, which involved the randomized presentation of annular visual gratings with varying drift rates, allowed us to assess the effect of increasing excitatory drive on grand-averaged GR parameters. However, this approach precluded analysis of how specific drift rates influenced trial-to-trial increases in gamma synchronization. For example, repetition-related changes in GR power might be more pronounced at higher drift rates, which elicit stronger excitation in the visual cortex. Future studies employing a modified experimental paradigm, such as presentation of specific drift rates in separate blocks, could address this question.

Another limitation is that our analysis was conducted at the sensor level rather than the cortical source level, which poses challenges, especially for interpreting evoked activity. Both the M80 and M180 components likely reflect the combined activity of multiple cortical regions, each potentially responding differently to stimulus repetition. Furthermore, we did not examine whether the habituation of ERF components varies as a function of drift rate, paralleling our approach to the analysis of single-trial GR power. Future studies utilizing source localization techniques and examining drift rate effects on ERF habituation could address these limitations.

Lastly, our VSS participants represented a heterogeneous group with various comorbidities. Due to the limited sample size, we were unable to reliably investigate the potential impact of comorbidities (e.g., anxiety, migraine with and without visual aura, etc.) on the MEG parameters analyzed. Understanding how these factors influence repetition-related gamma synchronization in VSS is another important avenue for future research.

In conclusion, this study provides the first independent confirmation of the previously observed stimulus repetition-related increase in MEG-recorded visual gamma oscillations, which has been associated with neuroplasticity and perceptual learning in healthy individuals. Our key finding was that this repetition-related gamma increase was unusually pronounced in patients with VSS, suggesting heightened Hebbian-type neuroplasticity in this disorder. At the same time, the normal modulation of gamma power and frequency by the strength of excitatory input to the visual cortex in VSS patients suggests that this enhanced plasticity is unlikely to result from an altered E-I ratio or a deficiency of inhibitory neurons involved in generation of gamma oscillations.

## Data availability

Data available on reasonable request.

## Acknowledgement

We warmly thank the patients and control group participants for participating in this study.

