## Supplementary table S1 for "Enhanced Neural Plasticity of the Primary Visual Cortex in Visual Snow Syndrome: Evidence from MEG Gamma Oscillations"

**Supplementary material**

Table S1. Spearman correlations between Gamma Response (GR) power regression coefficients and Heart Rate Variability (HRV) measures

| Parameter | Control + VSS | Control + VSS | |
| --- | --- | --- | --- |
|  | Rest (N = 48) | | Visual task (N = 52) |
| RMSSD | 0.2 | | 0.16 |
| SDNN | 0.05 | | -0.03 |
| pnn50 | 0.18 | | 0.18 |
| HF | 0.19 | | 0.15 |
| LF | -0.11 | | -0.15 |
| **HFnu** | **0.32*** | | **0.30*** |

* p < 0.05
